# Spatially Organized Tertiary Lymphoid Structures Emerge in Small Cell Lung Cancer and Associate with Improved Survival

**DOI:** 10.64898/2026.08.08.743663

**Authors:** Yingying Cao, Michael Nirula, Paul Mallory, Sarthak Sahoo, Kanak Parmar, Christopher A. Febres-Aldana, Anish Thomas

**Affiliations:** Developmental Therapeutics Branch, Center for Cancer Research, National Cancer Institute, National Institutes of Health, Bethesda, Maryland, USA; Imaging Mass Cytometry Laboratory, Frederick National Laboratory for Cancer Research, Leidos Biomedical Research, Inc., Frederick, MD, USA; Department of Bioengineering, Indian Institute of Science, Bengaluru, India; Laboratory of Pathology, Center for Cancer Research, National Cancer Institute, Bethesda, MD, 20892, USA

## Abstract

Tertiary lymphoid structures (TLS) are ectopic immune aggregates associated with improved prognosis and response to immunotherapy in multiple solid tumors. However, their presence, spatial organization, and functional relevance in small cell lung cancer (SCLC), a malignancy characterized by profound immune evasion, remain poorly understood. Using imaging mass cytometry (IMC) across 320 regions of interest spanning primary lung tumor, tumor-adjacent lung, liver and lymph node metastasis, complemented by Visium HD spatial transcriptomics, we characterized the cellular architecture and molecular programs of TLS-like niches in SCLC. TLS-like niches were identified in a subset of SCLC samples, predominantly primary lung tumor tissues and adjacent lung, spanning a continuum from loose lymphoid aggregates to compact follicle-like immune structures. Organized TLS-like niches contained CD20+ B-cell cores, closely associated with CD4+ and CD8A+ T cells, proliferating lymphocytes, HLA-DR+ antigen-presenting compartments, and αSMA+ stromal scaffolds, and were enriched for canonical TLS organizer signals (CXCL13, LTB, FDCSP). Patients with TLS-positive tumors demonstrated improved overall survival, and core TLS-associated transcriptional programs were associated with favorable survival in an independent bulk RNA-seq cohort. To our knowledge, this represents one of the first spatially resolved analyses of TLS-like immune architecture in SCLC, demonstrating that organized lymphoid immunity can emerge in this classically immune-evasive disease and is associated with improved survival.

## Introduction

Tertiary lymphoid structures (TLS) are ectopic, lymph node-like immune aggregates that arise in chronically inflamed tissues, including tumors ^1,2^. These structures recapitulate key features of secondary lymphoid organs, such as organized B cell follicles, T cell zones, and networks of antigen-presenting cells, enabling local antigen presentation and adaptive immune priming ^1,3^. Across multiple cancer types, the presence of TLS has been strongly associated with enhanced anti-tumor immunity, improved patient survival, and increased responsiveness to immunotherapy, particularly immune checkpoint blockade ^4–7^. Mechanistically, TLS are thought to serve as intratumoral hubs for coordinated immune activation, supporting B cell maturation, T cell activation, and sustained immune surveillance within the tumor microenvironment ^1,3^.

Despite their importance in other malignancies, the existence and functional relevance of TLS in small cell lung cancer (SCLC) remain poorly understood. SCLC is characterized by rapid progression, early metastasis, and a profoundly immunosuppressive microenvironment, with generally limited and heterogeneous responses to immunotherapy ^8–11^. Although TLS have been described in non-small cell lung cancer ^5,12^, evidence for organized lymphoid immunity in SCLC is sparse. In particular, the spatial organization, compositional features, and maturation states of TLS in SCLC have not been systematically characterized at single-cell resolution.

TLS are traditionally identified through histopathological assessment of lymphoid organization, including follicular organization and germinal center formation ^1,2,4^. However, translating these qualitative criteria to high-dimensional multiplexed imaging datasets remains challenging. Existing computational and image-based approaches have begun to address this limitation but often rely primarily on predefined marker combinations, morphology, or simple immune cell-density metrics ^4,13,14^, which may not fully distinguish spatially organized TLS from diffuse lymphoid aggregates. High-dimensional spatial profiling technologies, including imaging mass cytometry (IMC) and spatial transcriptomics platforms such as Visium HD, now provide an opportunity to interrogate immune architecture within intact tissues and to characterize TLS organization at single-cell proteomic and transcriptomic resolution.

Here, we used IMC across 320 regions of interest spanning primary tumors, adjacent lung tissue, liver metastases, and lymph node metastases, complemented by Visium HD spatial transcriptomics, to characterize TLS-like immune niches in SCLC. Our analyses demonstrate that spatially organized lymphoid structures can emerge within the SCLC microenvironment and exist along a continuum from loose immune aggregates to compact follicle-like niches enriched for canonical TLS programs. These structures exhibited coordinated B-cell, T-cell, antigen-presentation, and stromal organization and were associated with favorable clinical outcomes, supporting TLS architecture as a biologically and clinically relevant feature of the SCLC tumor microenvironment.

## Results

### Study cohort and IMC profiling

We assembled a tissue microarray (TMA) cohort of 320 regions of interest (ROIs) across seven TMAs derived from patients with pathologically confirmed SCLC. ROIs spanned four major anatomical compartments that formed the basis for site-level analyses: primary lung tumor (n = 147), adjacent lung tissue (n = 66), liver metastasis (n = 47), and lymph node metastasis (n = 27), with additional ROIs from adrenal (n = 7), bone (n = 5), and other metastatic sites (n = 21) included in cohort-wide but not site-level analyses (Fig. 1A; Supplementary Table S1). The cohort comprised 280 unique tissue samples. Overall survival data were available for 164 patients with tumor-site samples; median overall survival was 17.4 months, and the majority of observations (> 90%) represented death events, consistent with the aggressive clinical course of SCLC. Clinical covariates including age, sex, and pathological stage were available for a subset of the cohort and are summarized in Supplementary Table S1.

**Figure 1.**
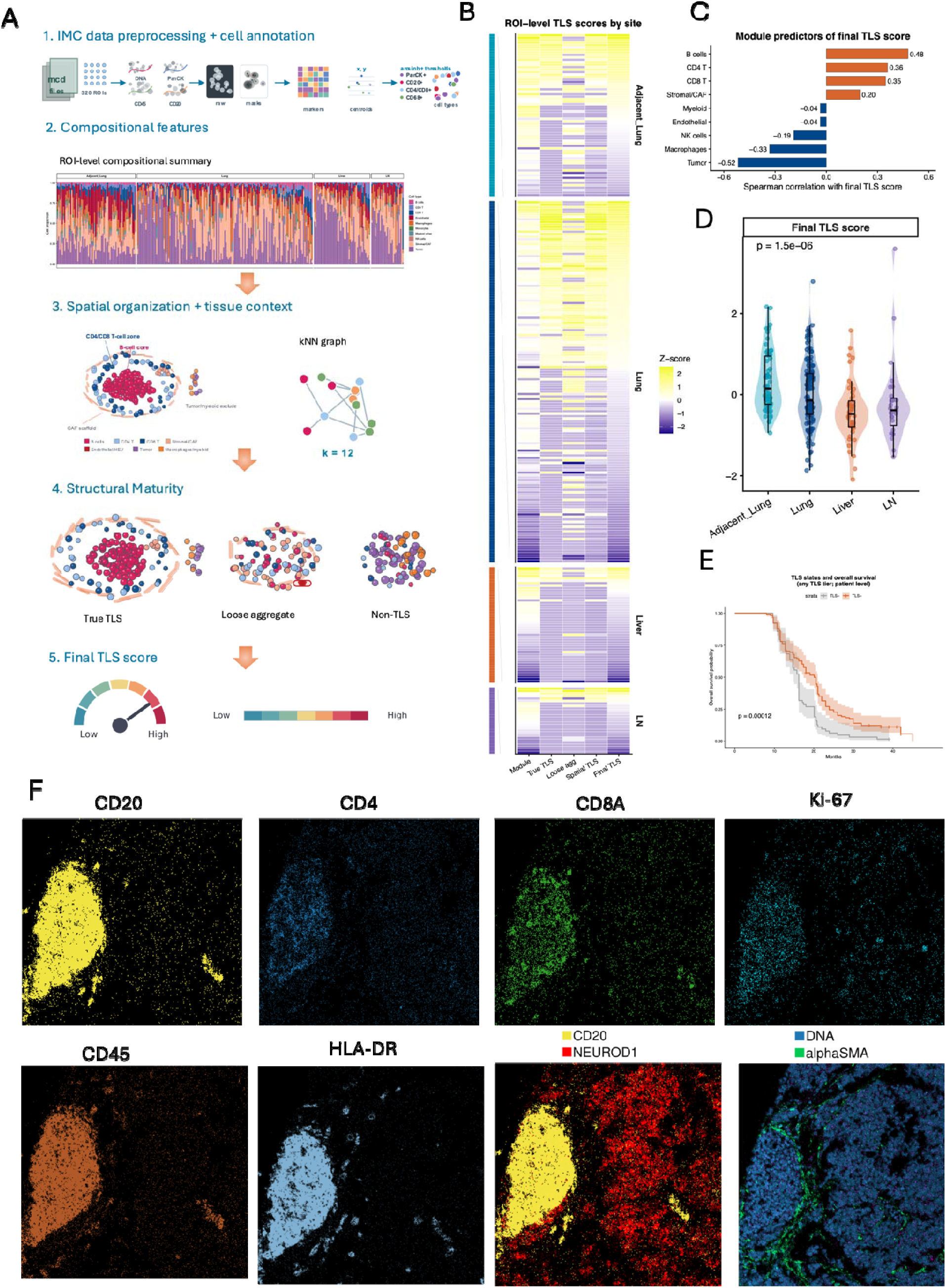
Spatial IMC analysis identifies TLS-like immune niches in SCLC and links organized TLS architecture to survival. **(A)** Overview of IMC workflow and spatial TLS analysis strategy. FFPE tissue microarray cores were stained with metal-conjugated antibodies, imaged by IMC, segmented into single cells, and annotated into major immune, stromal, and tumor cell populations. ROI-level TLS features were derived from five analytical layers: IMC preprocessing and cell annotation, cell-type compositional features, spatial immune organization and tissue context, structural maturity classification, and final integrated TLS scoring. Spatial immune organization was modeled using k-nearest-neighbor graphs, and immune clusters were classified as structured TLS-like niches, loose aggregates, or non-TLS immune clusters. **(B)** Heatmap of ROI-level TLS-associated features across anatomical sites. Rows represent ROIs grouped by tissue site, and columns represent TLS-related scores including compositional module score, true TLS score, loose aggregate score, spatial TLS score, and final integrated TLS score. Values are z-scored across ROIs. Adjacent lung and primary lung sites show higher TLS-associated organization than liver metastases and lymph node samples. **(C)** Cell-type module predictors of the final TLS score. Spearman correlations between ROI-level module features and the final TLS score are shown. Positive correlations are shown in orange and negative correlations in blue. B cells, CD4 T cells, CD8 T cells, and stromal/CAF features were positively associated with TLS score, whereas tumor and macrophage-related modules showed negative associations. **(D)** Distribution of final TLS scores by anatomical site. Violin and box plots show ROI-level final TLS scores across adjacent lung, primary lung tumor, liver metastasis, and lymph node samples. P value was calculated using a Kruskal-Wallis test. Adjacent lung and primary lung samples show higher and more variable TLS scores than metastatic sites. **(E)** Kaplan-Meier analysis of overall survival stratified by TLS status. Survival analysis was performed at the patient level using tumor-site samples, excluding adjacent lung from the prognostic model. Patients with TLS-positive tumors showed improved overall survival compared with TLS-negative patients. P value was calculated using a log-rank test. **(F)** Representative IMC images of a high-scoring TLS-like niche within a primary lung tumor sample. Single-marker and merged images show a compact CD20-positive B-cell aggregate adjacent to tumor and stromal compartments, including NEUROD1-positive tumor cells, nuclear DNA signal, and αSMA-positive stromal structures. These representative images illustrate the spatial organization underlying high TLS scores and support the distinction between structured TLS-like niches and diffuse immune infiltration.

All tissue sections were profiled using a 35-marker IMC panel encompassing immune lineage and activation markers (CD45, CD3, CD4, CD8a, CD20, CD68, Ki-67, HLA-DR, Granzyme B), myeloid and macrophage subset markers (CD163, TREM2, SPP1, MRC1, APOE), stromal and vascular markers (αSMA, CD31), tumor markers (Pan-Cytokeratin, NEUROD1), and additional microenvironment markers (Supplementary Table S2). Single-cell segmentation yielded 6,332,698 cells across 320 ROIs. Rule-based hierarchical phenotyping classified cells into major populations including tumor cells, B cells, CD4+ and CD8+ T cells, NK cells, macrophage subsets, cancer-associated fibroblasts, and endothelial cells, providing the cellular foundation for spatial TLS characterization across the SCLC tumor microenvironment.

### TLS-like immune niches are present in a subset of SCLC tumors

To characterize TLS-like immune organization in SCLC, we developed a multi-layer analytical framework that integrates compositional, spatial, and structural features at the ROI level, enabling systematic identification of TLS-like immune niches across the cohort. This framework operates on three complementary layers: a compositional layer that quantifies enrichment of lymphoid and stromal cell populations relative to suppressive and tumor compartments; a spatial layer that identifies local immune cell aggregation and tumor-immune organization using k-nearest-neighbor graphs and community detection; and a structural maturity layer that evaluates cluster-level features to distinguish organized TLS-like structures from loose immune aggregates. These layers are integrated into a unified TLS score that captures both the presence and biological quality of TLS-like structures. All analyses were performed using the full IMC dataset comprising 6,332,698 cells across 320 ROIs and 7 tissue microarrays, ensuring unbiased quantification of rare immune populations and spatial features (Fig. 1A).

Single-cell phenotyping identified major cell populations including tumor cells, B cells, CD4+ and CD8+ T cells, stromal and cancer-associated fibroblast (CAF) populations, and myeloid subsets. For each ROI, we computed a composite TLS score integrating three feature domains: compositional enrichment (relative lymphocyte abundance and diversity), spatial organization (neighborhood structure from k = 12 nearest-neighbor graphs), and tissue context (immune aggregate proximity to tumor boundaries). TLS scores varied substantially across anatomical sites (Fig. 1B, D). Adjacent lung tissue showed the highest TLS scores and the greatest frequency of structured immune aggregates, potentially reflecting both tumor-associated TLS and pre-existing pulmonary lymphoid organization, followed by primary lung tumors. In contrast, liver metastases and lymph node metastases exhibited significantly lower TLS scores (Kruskal-Wallis P = 1.5 × 10^-^□), consistent with more immunosuppressive or immune-excluded microenvironments at metastatic sites. At the cellular level, B cell and T cell abundance were the strongest positive predictors of the final TLS score (Spearman ρ = 0.48, 0.36, and 0.35 for B cells, CD4+ T cells, and CD8+ T cells, respectively), whereas tumor cell abundance was inversely correlated (ρ = −0.52) (Fig. 1C). These findings support a model in which coordinated lymphoid recruitment and stromal organization promote TLS formation, whereas tumor-dominant and macrophage-rich microenvironments are less permissive for organized lymphoid immunity.

Importantly, spatial analysis revealed that not all B-cell rich regions correspond to mature TLS-like structures. By incorporating spatial organization and local microenvironment features, we distinguished high-confidence true TLS clusters from looser immune aggregates. Organized TLS-like structures were characterized by high B-T cell co-localization, increased cluster density, and reduced tumor and macrophage infiltration. These structures were enriched in adjacent lung and present in a subset of primary tumor regions, whereas loose aggregates were more broadly distributed and less site-specific. This distinction demonstrates that immune infiltration alone is insufficient to define TLS and that spatially organized TLS-like niches represent a more biologically informative feature of the tumor microenvironment. Within each anatomical site, ROIs displayed a wide range of TLS scores, indicating substantial intratumoral heterogeneity. Rather than forming discrete categories, ROIs spanned a continuous spectrum from tumor-dominated, immune-poor regions through transitional mixed regions to highly organized TLS-rich niches. This gradient highlights the importance of ROI-level resolution in capturing tumor microenvironment diversity (Fig. 1B).

To assess the clinical relevance of TLS organization, we performed patient-level survival analysis in 164 patients with tumor-site samples and available overall survival data, excluding adjacent lung tissue from the prognostic model to avoid treating matched tumor-adjacent tissue as an independent tumor observation. TLS status was defined at the patient level: a patient was classified as TLS-positive if any of their tumor-site ROIs harbored definitive TLS (n = 37 TLS-positive; n = 127 TLS-negative). Patients with TLS-positive tumors showed significantly improved overall survival compared with TLS-negative patients (median OS 21.0 vs. 17.4 months; log-rank P = 0.00012; Fig. 1E), supporting TLS-like organization as a prognostically relevant feature in SCLC. Representative high-scoring ROIs from primary lung tumors showed compact, follicle-like CD20+ B-cell aggregates sharply demarcated from adjacent NEUROD1+ SCLC tumor nests, with the two compartments occupying mutually exclusive spatial domains (Fig. 1F). Within these structures, CD20+-cell cores were closely associated with intermingled CD4+ helper and CD8A+ cytotoxic T-cell populations, together forming pan-leukocyte (CD45+) aggregates with the characteristic compartmentalized architecture of organized lymphoid tissue. Ki-67 positivity within the aggregates indicated active lymphocyte proliferation, while strong HLA-DR expression confirmed local antigen-presentation activity consistent with functional TLS-like organization. The entire immune compartment was embedded within an αSMA+ fibroblastic stromal scaffold that delineated tumor-immune interfaces. Across cores, these TLS-like structures spanned a continuum from tightly organized, follicle-like CD20+ aggregates to more dispersed lymphoid clusters, consistent with varying degrees of TLS maturation within the SCLC microenvironment.

### Visium HD resolves molecular gradients surrounding TLS-like niches

To define the molecular architecture of TLS-like niches at higher spatial resolution, we performed cell-segmented Visium HD spatial transcriptomics on a representative TLS-containing primary lung tumor sample harboring three spatially discrete immune aggregates. Unsupervised clustering identified five major cell populations, including B cells, T cells, myofibroblastic CAFs, SPP1-positive tumor-associated macrophages, and tumor cells. These populations showed clear separation in UMAP space and distinct localization in tissue coordinates, with B- and T-cell–enriched aggregates embedded within a predominantly tumor- and myCAF-rich tissue landscape (Fig. 2A, B). High-resolution H&E imaging of the same tissue region further supported the presence of lymphoid aggregate-like structures at the corresponding locations (Fig. 2C).

**Figure 2.**
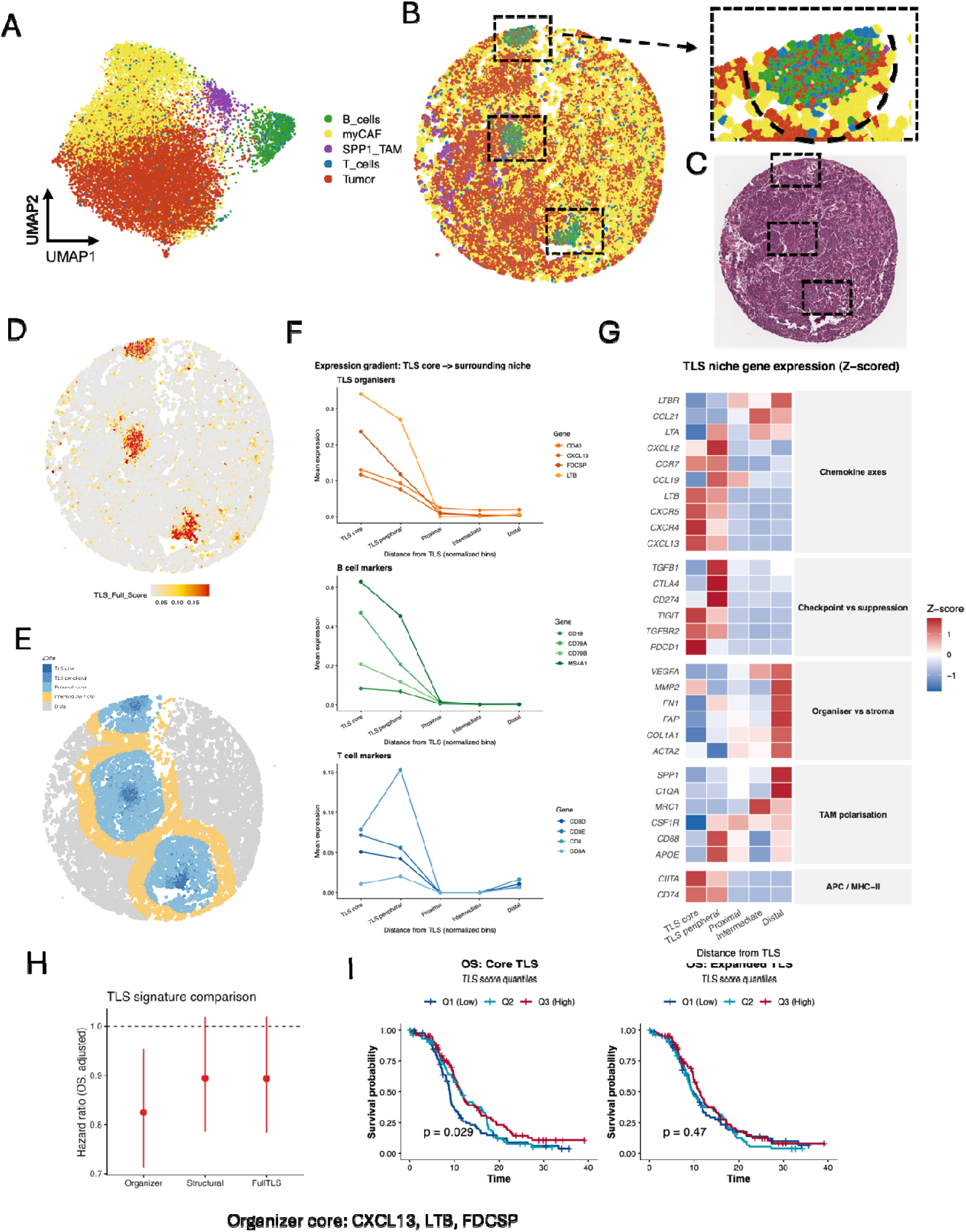
Visium HD spatial transcriptomics reveals radial molecular organization surrounding TLS-like niches in SCLC. **(A)** Cell-segmented Visium HD analysis of a representative TLS-containing SCLC tissue section. UMAP visualization of segmented cells identifies major annotated populations, including B cells, T cells, myofibroblastic cancer-associated fibroblasts (myCAF), SPP1-positive tumor-associated macrophages (SPP1_TAM), and tumor cells. **(B-C)** Spatial localization of annotated cell populations across the tissue section. TLS-like niches appear as spatially restricted B- and T-cell enriched aggregates embedded within a tumor- and myCAF-rich microenvironment. Dashed boxes indicate TLS-like regions, with corresponding magnified views and high-resolution H&E imaging of this representative TLS-containing SCLC tissue section **(C)**. **(D)** Spatial map of TLS score across the tissue section. Regions with high TLS score localize to discrete immune aggregates, highlighting three TLS-like niches within the sample. **(E)** Definition of TLS-centered spatial zones. TLS cores were identified by DBSCAN-based clustering of B and T cell positions. Cells were then assigned to concentric distance-based zones relative to the nearest TLS core: TLS core, TLS periphery, proximal niche, intermediate niche, and distal tissue. These zones were used to quantify radial changes in cell composition and gene expression. **(F)** Spatial gene expression gradients from TLS core to surrounding niche. Canonical TLS organizer genes, including CXCL13, LTB, and FDCSP, were enriched in the TLS core and decreased with increasing distance. B-cell markers showed strong core enrichment, while T-cell markers extended into the TLS periphery, indicating compartmentalized lymphoid organization. **(G)** Z-scored heatmap of TLS niche gene expression across spatial zones. Genes are grouped into functional programs, including chemokine axes, checkpoint and suppressive signaling, organizer and stromal programs, and tumor-associated macrophage polarization. TLS cores were enriched for lymphoid organizer and immune regulatory signals, whereas distal regions showed stronger stromal, extracellular matrix, angiogenic, and macrophage-associated programs. **(H)** Comparison of TLS-associated transcriptional signatures. Adjusted hazard ratios are shown for organizer, structural, and full TLS signatures, indicating the prognostic relationship between TLS-related molecular programs and overall survival. **(I)** Kaplan-Meier analysis of overall survival stratified by core TLS score and expanded TLS score quantiles. Higher core TLS signal was associated with improved survival, supporting the clinical relevance of compact TLS-centered immune organization. Expanded or diffuse TLS-associated signal was not significantly associated with survival, indicating that compact TLS core organization, rather than broad lymphoid expansion alone, captures the clinically relevant TLS feature.

To examine spatial organization surrounding TLS-like niches, we defined TLS-centered spatial zones using DBSCAN-based clustering of B- and T-cell positions, followed by distance-based assignment of all segmented cells relative to the nearest TLS core. This generated five concentric compartments: TLS core, TLS periphery, proximal niche, intermediate niche, and distal tissue (Fig. 2D, E). Cell-type composition changed markedly across these zones. B and T cells were enriched in the TLS core, myCAF and tumor cells became increasingly dominant with distance from the TLS core. These patterns indicate that TLS-like structures are organized as spatially restricted immune niches rather than diffuse lymphoid infiltrates (Fig. 2B, F).

Spatial transcriptomic analysis revealed coordinated molecular gradients extending from the TLS core into the surrounding tumor microenvironment (Fig. 2F, G). Canonical TLS organizer genes, including CXCL13, LTB, and FDCSP, were highest within the TLS core and decreased sharply with distance. The CXCL12-CXCR4 axis showed a similar core-enriched pattern, consistent with active lymphocyte retention and follicular organization. B-cell markers (MS4A1, CD19, CD79A, CD79B) showed a similar core-enriched pattern, confirming the lymphoid identity of these structures. T-cell markers (CD3D, CD3E, CD4, CD8A) were enriched in the TLS core and periphery, consistent with compartmentalized lymphoid organization. Immune checkpoint and regulatory genes, including PDCD1, CTLA4, CD274, TIGIT, TGFB1, and TGFBR2, were similarly concentrated within TLS-centered zones, consistent with localized T-cell activation accompanied by feedback regulation rather than diffuse immune exhaustion.

In contrast, stromal and pro-tumorigenic programs showed reciprocal spatial enrichment outside the TLS niche. Extracellular matrix and stromal remodeling genes, including COL1A1, ACTA2, FN1, FAP, and MMP2, together with the angiogenic factor VEGFA, were enriched in distal myCAF- and tumor-dominated regions. The macrophage compartment was spatially zoned by phenotype: CD68+ and APOE+ monocyte-derived macrophages localized at the TLS periphery, MRC1+ M2-skewed macrophages occupied intermediate zones, and SPP1+ and C1QA+ pro-tumorigenic macrophages dominated the distal stroma, mutually excluded from the TLS niche. A Z-scored heatmap across TLS-centered zones confirmed this radial organization, with TLS-core immune organizer, antigen-presentation, and regulated checkpoint programs spatially opposing distally enriched stromal, angiogenic, and SPP1-driven myeloid programs (Fig. 2G).

To assess whether TLS-associated transcriptional programs carried prognostic information beyond our IMC cohort, we analyzed bulk RNA-seq data from the IMpower133 trial ^15^, a phase III randomized study of atezolizumab plus carboplatin-etoposide versus placebo plus carboplatin-etoposide in 271 patients with extensive-stage SCLC that compared atezolizumab plus carboplatin-etoposide with placebo plus carboplatin-etoposide. We evaluated two TLS-associated signatures by ssGSEA across the full intention-to-treat population: a core TLS organizer signature (CXCL13, LTB, FDCSP) and an expanded TLS signature incorporating broader lymphoid, antigen-presentation, and immune activation genes. Patients were stratified into tertile groups by signature score. The core TLS organizer signature was significantly associated with improved overall survival, with a dose-response pattern across score tertiles (log-rank P = 0.029; Fig. 2H, I). This association remained significant after adjustment for treatment arm, adjusted for treatment assignment, including atezolizumab plus chemotherapy versus placebo plus chemotherapy, transcriptional subtype, ECOG performance status, brain and liver metastasis status, and LDH level in a multivariable Cox model (Fig. 2H). In contrast, the expanded TLS signature was not significantly associated with overall survival (P = 0.47), suggesting that the clinically relevant TLS signal in SCLC is concentrated in compact TLS organizer programs rather than in diffuse lymphoid expansion. These results indicate that core TLS-associated molecular programs have prognostic value in SCLC independent of established clinical covariates and treatment assignment.

## Discussion

In this study, we performed a spatially resolved, multi-modal characterization of TLS-like immune niches in SCLC, a tumor type widely regarded as immunologically cold and in which organized lymphoid immunity has been minimally explored ^1,9,16^. By integrating imaging mass cytometry across 320 ROIs with Visium HD spatial transcriptomics, we demonstrate that TLS-like structures can emerge within the SCLC microenvironment, that they exist along a continuum of structural maturation, and that their presence is associated with improved overall survival. These findings suggest that the microenvironmental organization required for coordinated anti-tumor immunity is preserved in a subset of SCLC ^8,17^. Prior single-cell and spatial studies have shown that SCLC is often immunosuppressive but also spatially heterogeneous, with specific immune niches linked to clinical outcome ^18,19^. Our findings extend this concept by identifying spatially organized TLS-like lymphoid niches as a biologically and clinically relevant form of adaptive immune organization in SCLC, consistent with broader evidence linking TLS to favorable anti-tumor immunity across multiple cancer types ^7^.

A central insight from this work is that TLS in SCLC is not a binary phenomenon. Our multi-layer scoring framework, which integrates compositional enrichment, spatial clustering, and architectural maturity, revealed that immune aggregates in SCLC span a broad spectrum from loose, partially organized B-cell rich clusters to compact, spatially structured follicle-like niches with coordinated B- and T-cell organization, proliferative activity, antigen-presentation programs, and stromal scaffolding. This continuum mirrors observations in other solid tumor types, where TLS maturation state has been linked to differential prognostic and immunomodulatory significance. Critically, our analyses demonstrate that immune cell abundance alone is insufficient to define TLS: many B-cell rich regions lacked the spatial organization and microenvironmental context that distinguish functional TLS from nonspecific immune infiltration. This distinction has practical implications for biomarker development, as approaches that rely solely on cell-type proportions or marker thresholds may overestimate the prevalence of functionally relevant TLS.

The spatial distribution of TLS-like niches across anatomical sites was markedly heterogeneous. Adjacent lung tissue consistently harbored the highest TLS scores and primary lung tumors contained TLS-like niches in only a subset of ROIs. Liver metastases and lymph node metastases showed significantly lower TLS scores, consistent with more immunosuppressive or immune-excluded microenvironments. The enrichment of TLS-like structures within adjacent lung tissue raises the possibility that some aggregates may represent pre-existing bronchus-associated lymphoid tissue or chronic inflammatory lymphoid organization rather than exclusively tumor-induced TLS. However, organized TLS-like niches were also identified within primary tumor regions, where they formed clear spatial interfaces with adjacent neuroendocrine tumor nests and were associated with favorable clinical outcome, supporting their biological relevance within the SCLC tumor microenvironment. More broadly, these findings suggest that pulmonary lymphoid organization and tumor-associated TLS may exist along a continuum rather than as strictly distinct entities.

Visium HD spatial transcriptomics provided a high-resolution view of the molecular architecture surrounding TLS-like niches. The concentric zone analysis revealed sharply organized gradients: canonical TLS organizer genes (CXCL13, LTB, FDCSP) and lymphoid markers peaked at the TLS core and decayed rapidly with distance, stromal programs (COL1A1, ACTA2, FN1) showed reciprocal enrichment in the distal microenvironment. This spatial reciprocity defines a functional boundary between immune-permissive and immune-suppressive territories within the same tissue section. Antigen-presentation programs, including the MHC class II master regulator CIITA and the invariant chain CD74, was similarly enriched within the TLS niche, and was directly corroborated at the protein level by strong HLA-DR signal within CD20+ aggregates in IMC, supporting the interpretation that these structures function as localized sites of immune activation and antigen presentation. The concentration of immune checkpoint molecules (PDCD1, CTLA4, TIGIT) at the TLS core further suggests that these structures harbor activated T cells engaged in antigen recognition, raising the possibility that TLS serve as sites of active but potentially constrained immune responses in SCLC.

The prognostic association between TLS-like oragnization and overall survival in our cohort parallels findings in non-small cell lung cancer, pancreatic cancer, and other solid tumors ^4–6^, and extends this relationship to SCLC for the first time using a spatially defined scoring framework. Notably, compact core TLS-associated transcriptional programs demonstrated stronger prognostic association than broader diffuse lymphoid signatures, suggesting that spatially organized immune architecture, rather than generalized immune infiltration alone, captures the clinically relevant TLS signal in SCLC. While these analyses were performed in relatively modest cohorts and should therefore be interpreted cautiously, the consistency between IMC-based spatial findings and independent bulk transcriptomic survival analyses supports the biological relevance of TLS-associated immune organization in this disease context. Future studies in larger independent cohorts will be important to validate TLS as a prognostic biomarker and to determine whether TLS status predicts responsiveness to immune checkpoint blockade or other immunotherapeutic strategies in SCLC.

Several limitations of this study should be acknowledged. First, although IMC provides high-dimensional protein profiling at single-cell resolution, tissue microarray sampling inherently captures only limited tissue areas and may not fully represent intratumoral spatial heterogeneity. Second, our Visium HD analysis was performed on a single representative TLS-containing sample, and while it provides a detailed molecular portrait, broader spatial transcriptomic profiling across additional samples and sites will be necessary to assess the generalizability of the molecular gradients described. Third, although our spatial analysis integrated compositional, organizational, and structural features of TLS-like niches, classification of TLS maturity remains dependent on predefined thresholds and spatial criteria that may benefit from further refinement in larger datasets. Finally, the functional consequences of TLS formation in SCLC cannot be fully resolved from observational spatial analyses alone, and future mechanistic studies will be required to determine whether these structures actively support productive anti-tumor immunity or reflect partially constrained immune activation within the SCLC microenvironment.

In summary, this study provides a spatially resolved characterization of TLS-like immune niches in SCLC, and demonstrates that organized lymphoid immunity can emerge even within this highly immune-evasive malignancy. By integrating multiplex spatial proteomics and spatial transcriptomics, we identify coordinated cellular and molecular programs that define TLS architecture and its interface with the surrounding tumor microenvironment. These findings expand the landscape of immune organization in SCLC and offer a foundation for future investigations into the therapeutic relevance of TLS in this challenging malignancy.

## Methods

### TLS quantification and spatial analysis

For each ROI, cell-type proportions were calculated by dividing the number of cells assigned to each cell type by the total number of cells in that ROI. Proportions were aggregated into biologically meaningful modules: lymphoid modules (B cells, CD4+ T cells, CD8+ T cells, NK cells), support modules (stromal/CAF and endothelial cells), and suppressive modules (macrophages, myeloid cells, tumor cells). Each module was standardized across ROIs using z-score transformation. A TLS module score was defined as the mean of the positive module z-scores (lymphoid and support modules) minus the mean of the negative module z-scores (suppressive modules), capturing the degree of lymphoid enrichment relative to suppressive and tumor compartments.

To identify candidate TLS regions, spatial clustering was performed on immune cells (B cells, CD4+ T cells, CD8+ T cells, CD8+ cytotoxic T cells, and NK cells) within each ROI. A k-nearest-neighbor graph (k = 12) was constructed using cell centroid coordinates, and communities were detected using the Louvain algorithm. Clusters with fewer than 15 cells were excluded. For each immune cluster, we computed cell composition (B, T, NK fractions), spatial density (cells per convex hull area), and local tissue context (tumor, macrophage, and stromal proportions in a surrounding neighborhood).

Each immune cluster was assigned a TLS maturity score integrating lymphoid composition (B/T/NK enrichment), cluster density (compactness), stromal and endothelial support, and absence of tumor and macrophage dominance. Clusters were classified into three categories: true TLS (high lymphoid content, balanced B-T composition, low tumor and macrophage infiltration, high maturity score), loose aggregates (partial lymphoid enrichment without full spatial organization or supportive microenvironment), and non-TLS clusters (remaining immune clusters).

For each ROI, spatial TLS features were summarized as the number of TLS clusters, maximum TLS maturity score, number and size of loose aggregates, and a combined spatial TLS score defined as TLS_spatial_score = TLS_true_score + 0.5 × TLS_loose_score. A final composite ROI-level TLS score was computed by averaging the z-scored TLS module score and the z-scored TLS spatial score. ROIs in the top 10% of the TLS score distribution were classified as TLS-high.

Differences in TLS-related scores across anatomical sites were assessed using Kruskal-Wallis tests. Associations between TLS score and cell-type modules were evaluated using Spearman correlation with multiple testing correction using the Benjamini-Hochberg method. Overall survival was compared between TLS-positive and TLS-negative patient groups using Kaplan-Meier analysis with log-rank tests. Correlations between the final TLS score and individual TLS module components and raw cell-type composition features were computed using Spearman rank correlation to identify features most strongly associated with TLS formation. Site-level summaries were used for descriptive biological comparisons and included adjacent lung, primary lung tumor, liver metastasis, lymph nodes. Because adjacent lung and tumor samples were frequently paired within the same patient, adjacent lung was excluded from the prognostic survival analyses. Survival models were performed at the patient level using tumor-site samples only, with TLS status summarized across available tumor sites per patient. Overall survival was compared using Kaplan–Meier analysis with log-rank testing, and hazard ratios were estimated using Cox proportional hazards models.

### Analysis of Visium HD spatial transcriptomics

Visium HD spatial transcriptomics was performed on a representative TLS-containing SCLC sample. High-resolution histology image was used to segment individual cells within the Visium HD tissue section. Transcript counts were assigned to segmented cells according to their spatial overlap with Visium HD capture bins, generating a cell-by-gene expression matrix with corresponding spatial coordinates for each cell. Cells with low transcript counts, low detected gene numbers, or poor segmentation quality were excluded. Gene expression values were normalized and log-transformed before dimensionality reduction and clustering. Unsupervised clustering of spatially resolved transcriptomic profiles identified five major cell populations: B cells, T cells, myofibroblastic CAFs (myCAF), SPP1+ tumor-associated macrophages (SPP1_TAM), and tumor cells. Concentric spatial zones were defined around TLS cores using DBSCAN-based clustering of B and T cell positions, followed by distance-based stratification of all cells relative to the nearest TLS core into five zones: TLS core, TLS peripheral, proximal niche, intermediate niche, and distal. Gene expression was analyzed across these zones to identify spatial gradients in TLS-associated, immune checkpoint, immunosuppressive, and stromal gene programs.

For survival analysis, gene expression and clinical annotation data were obtained from the Barzin et al. ^15^ bulk RNA-seq cohort comprising 271 patients with SCLC. Transcript per million (TPM) values were log2-transformed [log2(TPM + 1)] prior to downstream analysis. TLS-associated transcriptional signatures were evaluated using single-sample gene set enrichment analysis (ssGSEA) implemented in the GSVA framework. Two predefined signatures were analyzed: (i) a core TLS organizer signature consisting of canonical lymphoid organizer genes (CXCL13, LTB, FDCSP), and (ii) an expanded TLS signature incorporating broader lymphoid, antigen-presentation, and immune activation genes. Patient-level TLS scores were then integrated with clinical metadata, including OS, treatment status, transcriptional subtype, ECOG performance status, metastatic burden, and LDH levels. For survival analyses, patients were stratified into quantiles based on TLS signature scores. Kaplan-Meier survival curves were generated to compare OS across TLS score groups, with statistical significance assessed using two-sided log-rank tests. Cox proportional hazards models were used to estimate hazard ratios (HRs) for TLS-associated signatures. Core TLS and expanded TLS signatures were evaluated separately to determine whether compact, organizer-driven TLS programs or broader lymphoid transcriptional signatures were more strongly associated with clinical outcomes.

## Notes

### Competing Interest Statement

The authors have declared no competing interest.

### Summary of Updates

The author order and corresponding author designations have been updated to reflect the authors' contributions to the revised manuscript. No changes were made to the scientific content of the manuscript.

## References

1. Schumacher, T. N. & Thommen, D. S. Tertiary lymphoid structures in cancer. Science (1979). 375, (2022).

2. Pitzalis, C., Jones, G. W., Bombardieri, M. & Jones, S. A. Ectopic lymphoid-like structures in infection, cancer and autoimmunity. Nature Reviews Immunology 2014 14:7 14, 447–462 (2014).

3. Deng, S. et al. Tertiary lymphoid structures in cancer: spatiotemporal heterogeneity, immune orchestration, and translational opportunities. Journal of Hematology & Oncology 2025 18:1 18, 97-(2025).

4. Sautès-Fridman, C., Petitprez, F., Calderaro, J. & Fridman, W. H. Tertiary lymphoid structures in the era of cancer immunotherapy. Nature Reviews Cancer 2019 19:6 19, 307–325 (2019).

5. Dieu-Nosjean, M. C. et al. Long-term survival for patients with non-small-cell lung cancer with intratumoral lymphoid structures. Journal of Clinical Oncology 26, 4410–4417 (2008).

6. Hiraoka, N. et al. Intratumoral tertiary lymphoid organ is a favourable prognosticator in patients with pancreatic cancer. British Journal of Cancer 2015 112:11 112, 1782–1790 (2015).

7. Vanhersecke, L. et al. Mature tertiary lymphoid structures predict immune checkpoint inhibitor efficacy in solid tumors independently of PD-L1 expression. Nature Cancer 2021 2:8 2, 794–802 (2021).

8. Thomas, A., Mohindroo, C. & Giaccone, G. Advancing therapeutics in small-cell lung cancer. Nature Cancer 2025 6:6 6, 938–953 (2025).

9. Kim, S. Y., Park, H. S. & Chiang, A. C. Small Cell Lung Cancer: A Review. JAMA 333, 1906–1917 (2025).

10. Poirier, J. T. et al. New Approaches to SCLC Therapy: From the Laboratory to the Clinic. Journal of Thoracic Oncology 15, 520–540 (2020).

11. Remon, J. et al. Small cell lung cancer: a slightly less orphan disease after immunotherapy. Annals of Oncology 32, 698–709 (2021).

12. Elfving, H. et al. Spatial distribution of tertiary lymphoid structures in the molecular and clinical context of non-small cell lung cancer. Cellular Oncology 2025 48:3 48, 801–813 (2025).

13. Li, Z. et al. Development and Validation of a Machine Learning Model for Detection and Classification of Tertiary Lymphoid Structures in Gastrointestinal Cancers. *JAMA Netw*. Open 6, e2252553–e2252553 (2023).

14. Wang, Y. et al. Computerized tertiary lymphoid structures density on H&E-images is a prognostic biomarker in resectable lung adenocarcinoma. iScience 26, (2023).

15. Nabet, B. Y. et al. Immune heterogeneity in small-cell lung cancer and vulnerability to immune checkpoint blockade. Cancer Cell 42, 429–443.e4 (2024).

16. Gazdar, A. F., Bunn, P. A. & Minna, J. D. Small-cell lung cancer: What we know, what we need to know and the path forward. Nat. Rev. Cancer 17, 725–737 (2017).

17. Qin, K., Gay, C. M., Byers, L. A. & Zhang, J. The current and emerging immunotherapy paradigm in small-cell lung cancer. Nature Cancer 2025 6:6 6, 954–966 (2025).

18. Jin, Y. et al. Single-cell and spatial proteo-transcriptomic profiling reveals immune infiltration heterogeneity associated with neuroendocrine features in small cell lung cancer. Cell Discov. 10, 93-(2024).

19. Zhang, Z. et al. Single-cell spatial transcriptomics reveals tumor microenvironment heterogeneity in primary and lymph node-metastatic small cell lung cancer. Cell Rep. Med. 7, 102713 (2026).

